# Enrichment of charge-absent regions in phase separated proteins

**DOI:** 10.1101/2022.11.05.515309

**Authors:** Sonia T. Nicolaou, Chandra S. Verma, Jim Warwicker

## Abstract

Many studies focus on the relationship between protein charge and liquid-liquid phase separation (LLPS), generally finding that a large degree of charge neutralisation is involved for condensate formation. Here, sequences within human proteins that lack the charge-bearing residues Asp, Glu, Lys, and Arg (termed charge-absent) are analysed alongside annotation for involvement in LLPS. Scaffold proteins, central to condensate formation, on average possess longer charge-absent regions than those not key for LLPS. Charge-absent regions tend to have relatively high hydropathy scores. Overall, they are enriched in Ala, Gly, Pro, and Ser with more specific groupings evident when the subset is clustered by amino acid composition. For several proteins, segments with charge-absent regions have been identified as modulators of LLPS. It is hypothesised that for at least some of the charge-absent regions, a lack of charged group desolvation energy, together with a relatively hydrophobic sequence composition, may facilitate condensation through homomeric interactions. If this is the case, it should be relatively easy to modulate through incorporation of charge through engineering, potentially including pH-sensing.

## Introduction

Cells organise their densely packed chemical space into compartments (organelles) as a way of regulating complex biochemical reactions. Typically, cellular components are encapsulated in membranes. Alternatively, these cellular components form membraneless organelles (MLOs), through liquid-liquid phase separation (LLPS), such as stress granules, Cajal bodies, P-bodies, PML bodies, and nuclear speckles. Recent studies have linked formation of MLOs to intrinsically disordered proteins (IDPs) [1], proteins that lack distinct secondary or tertiary structures and instead exist as dynamic conformational ensembles [2–6]. IDPs are composed of amino acid (AA) sequences with lower complexity since they are enriched in “disorder promoting” residues such as Ala, Arg, Gly, Gln, Ser, Glu, Lys and Pro and are depleted in hydrophobic and aromatic residues [4, 7, 8]. Low complexity domains (LCDs) are sometimes involved in LLPS [9–11]. Some of these low complexity sequences are called prionlike domains (PLDs) due to sequence similarities with yeast prion proteins [12]. The high net charge and low mean hydrophobicity of IDP sequences are an important contributor to their lack of compact structure, due to charge-charge repulsion [13]. The charge-hydrophobicity relationship of IDPs is clearly demonstrated in the charge-hydropathy plot, also known as the Uversky plot [5, 14]. Several disorder prediction servers utilise this principle to describe IDPs [15–17]. A diagram-of-states to classify the predicted conformational properties of IDPs based on their charge-hydropathy relationship was initially created by Mao and co-workers and was later improved by Das and Pappu [18, 19]. The two-dimensional (2D) diagram-of-states has fraction of positively charged residues on the x-axis (f+) and fraction of negatively charged residues (f-) on the y-axis and is divided into 5 regions based on their fraction of charged residues (FCR) and net charge per residue (NCPR) scores [19, 20].

IDPs adopt an ensemble of conformations, and due to their flexible nature, exposed AA sequence, and binding promiscuity, have an increased transient interaction network thought to enhance their phase separation properties [21–24]. Multivalent interactions between IDPs, IDPs and folded proteins, IDPs and nucleic acids, and nucleic acids-nucleic acids, including hydrogen bond, electrostatic, cation-π, π-π, dipole-dipole and hydrophobic interactions are the driving force of LLPS [25], along with modulations in the solution/environment of the system such as temperature, hydrostatic pressure, pH and ionic strength [26–28]. Under physiologic conditions, some IDPs undergo LLPS as a way of regulating important biological processes. However, changes in the sequence dependent phase behaviour of IDPs such as mutations and post-translational modifications (PTMs), can sometimes lead to the formation of pathological aggregates [21, 29]. Protein aggregation is the hallmark of many neurodegenerative diseases such as amyotrophic lateral sclerosis (ALS), frontotemporal dementia (FTD), Alzheimer’s disease (AD), and Parkinson’s (PD). TAR DNA-binding protein 43 (TDP-43), fused in sarcoma (FUS), tau, and α-synuclein are examples of IDPs found in patients with ALS, FTD, AD, and PD, respectively [29–34]. Under physiologic conditions, these proteins exist in a dispersed state and can undergo reversible transition to phase separated liquid droplets. However, under stress they form irreversible pathological aggregates.

While recent studies focused on the importance of charge in IDPs and its implications in LLPs, IDPs with regions of little or no charge also exist in the human proteome, such as in the carboxy-terminus of TDP-43. TDP-43 binds to DNA and RNA through two RNA recognition motifs (RRM1 and RRM2) [35]. It also contains a disordered, Gly-rich, low complexity (PLD) carboxy-terminus. This disordered terminal region contains a short, conserved region (CR) between residues 316-346, and can promote phase separation of TDP-43 into stress granules and cytosolic aggregates [12, 36]. The CR lacks charged AAs, but is enriched in polar ones (Gln, Asn, Ser and Gly) and contains hydrophobic residues Tyr, Phe and ten evolutionarily conserved Met residues [12, 37]. The CR forms an α-helical structure that is stabilised by intermolecular contacts between CR helices of adjacent TDP-43 molecules and is associated with condensation [36, 38].

The function of IDPs can generally be established from their amino acid sequence, often enriched in charged and polar residues. However, intrinsically disordered regions with very little or no residues expected to be charged at neutral pH pose the question of why such regions exist in the human proteome and if they are functionally significant. One in five AAs are charged (Asp, Glu, Lys, Arg) and therefore it is unlikely that long polypeptide segments without charge would exist by chance. Studies by Bremer et al. suggest that electroneutral NCPR values promote LLPS, whereas larger NCPR values (unbalanced charges) suppress LLPS [39]. The diagram-of-states [19] uses f+, f-, FCR and NCPR to describe the conformational preferences of IDPs. The current study focuses on the (0,0) point of the diagram to investigate regions of proteins with zero fraction of both positively and negatively charged residues (termed charge-absent regions). It is found that the critical scaffold proteins in phase separated condensates are enriched in charge-absent regions, and that these regions are as hydrophobic as structured proteins despite mostly being disordered and depleted in larger hydrophobic amino acids.

## Methods

### Dataset

The human proteome, with one protein sequence per gene (proteome ID UP000005640, 20588 sequences) was acquired from the UniProt database [40]. Proteins with transmembrane (TM) domains were excluded from the dataset since the study is focused on aqueous soluble proteins. Proteins were assumed to contain TM segments if their hydropathy score was greater than 1.6 on the Kyte & Doolittle [41] scale, for any 21 AA segment [42], which would be representative of a typical TM helix. After this filter, 623 proteins containing 51 AA stretches of zero charge (charge-absent, no K/R/D/E residues) were identified in the human proteome.

Prediction of intrinsically disordered segments was made with IUpred2A [43, 44]. Any single amino acid with score > 0.5 is predicted to be disordered. The AA composition of each 51 AA charge-absent segment was found using the web tool iFeature [45]. Proteins involved in LLPS were obtained from the DrLLPS database [46] and were matched with the subset of proteins containing charge-absent regions of at least 51 amino acids. The proteins were then divided into 3 categories: scaffold, client, and regulator.

The dataset for PrLDs used was acquired from March et al. [47]. Gene Ontology (GO) analysis for a given set of genes was performed using Princeton GO term finder [48–50], and Revigo [51] was used to visualise the results. An additional dataset for benchmarking hydrophobicity of the charge-absent subset was obtained from previous work [52], with human protein structures from the Protein Data Bank (PDB) [53], non-redundant at 25% sequence identity.

### Clustering

The AA compositions of the 623 51 AA charge-absent sequences (one per protein) were normalised by calculating their z-scores i.e. the number of standard deviations away from the mean (over the complete subset) for the composition of each AA type in each 51 AA window. Agglomerative hierarchical clustering of the normalised AA compositions of the sequences was performed using the Ward variance minimization algorithm, and the Euclidean distance function as the distance metric between vectors. During agglomerative hierarchical clustering each object (protein sequence) constitutes its own cluster, and then the clusters closest together (clusters with the shortest Euclidean distance between them) are successively merged [54–56]. The resulting output is a dendrogram with the Euclidean distance between clusters on one axis and the clusters on the other. SciPy was used to create the dendrogram and colour-coded each group of nodes whose linkage was less than 70% of the maximum linkage. The dendrogram was then visually inspected based on the SciPy colour-coding and 11 clusters were identified [57].

## Results and discussion

### Scaffold LLPS proteins are enriched in charge-absent regions

Previous charge distribution analysis showed that IDRs of proteins involved in LLPS are biased towards neutral net charge [17]. Human proteins in our dataset were divided into scaffold, client, and regulator proteins, or unannotated (null), based on DrLLPS, an online database that lists proteins involved in LLPS [46]. Scaffold proteins are directly involved in LLPS, client proteins are recruited into MLOs but are not essential for LLPS, and regulators modulate LLPS but are not localised in the droplets [58]. The null subset is that part of the human proteome not annotated in DrLLPS, and without predicted TM segments. Cumulative plots of the maximum charge-absent lengths were made for proteins with the associations scaffold, regulator, client in DrLLPS, as well as other human proteins (lacking a predicted TM segment and labelled as null in Fig 1). Interestingly the regulator, client, and null plots almost overlay, whilst that of proteins labelled as scaffold is enriched in longer charge-absent regions. The maximum separation, in the cumulative plots, between scaffold proteins and the other subsets is around a charge-absent window of 50 amino acids (Fig 1). It is believed that net charge is an important regulator of IDR conformation and global dimensions in solution [5, 17, 19, 20, 59, 60] and this observation is consistent with a model in which condensation of some human scaffold proteins is aided by relatively long intrinsically disordered regions bearing zero charge at neutral pH. For more detailed study, a length of contiguous charge-absent sequence was set to 51 amino acids. This is a balance of seeking a value toward the upper end charge-absent region length and maintaining a difference between scaffold and other subsets (Fig 1).

**Fig 1.**
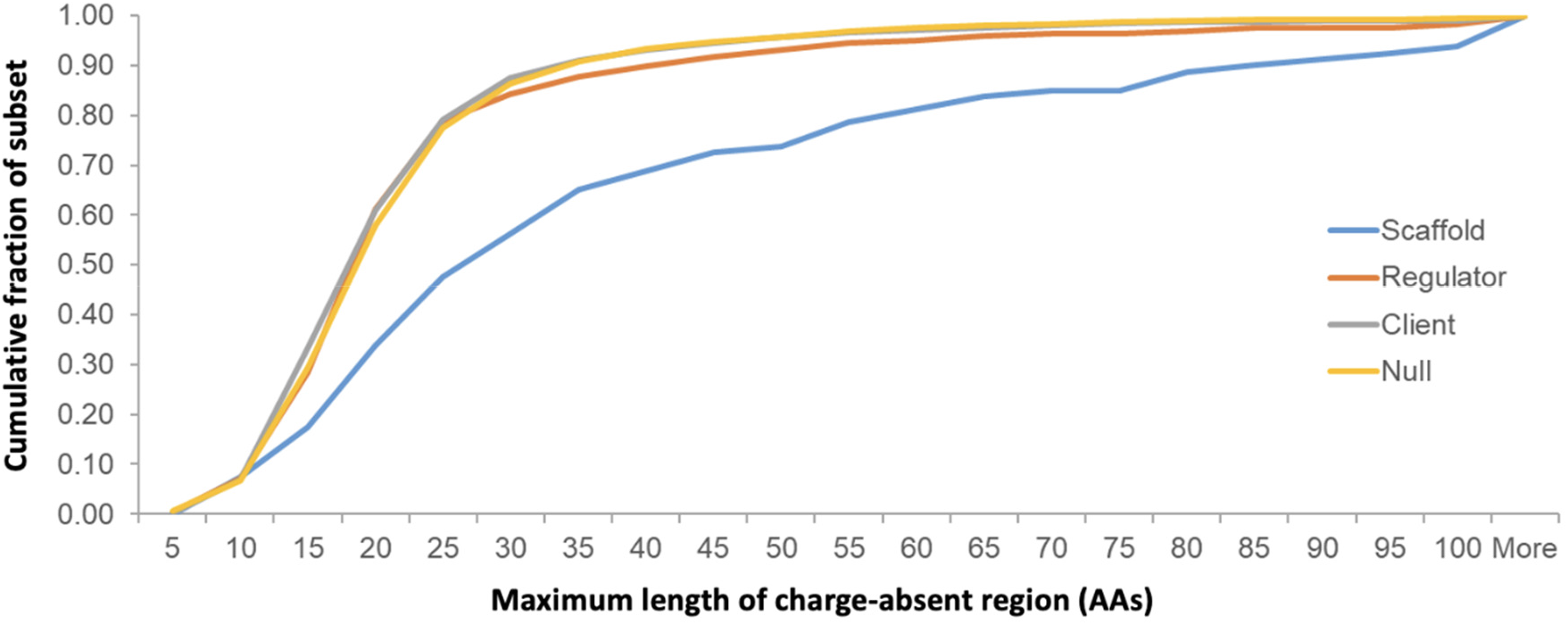
Scaffold proteins are enriched in longer charge-absent regions. Subsets of human proteins, according to annotation in DrLLPS (client N=80, regulator N=278, client N=2651), are displayed in cumulative distributions of maximum length of charge-absent region in each protein. The null subset covers human proteins without DrLLPS annotation (and lacking a predicted TM segment, N=10778). The plots for regulator, client, and null subsets almost overlay.

The first charge-absent segment, for each of the 623 proteins with at least one 51 AA charge-absent segment was submitted to the IUpred2A disorder predictor [43, 44]. The average IUPred2A score for the 623 51 charge-absent segments was 0.532, with 52.5% of amino acids having disorder prediction (scores >= 0.5). Moreover, 482 out of the 623 51 regions had at least one AA with score >= 0.5. The majority of the charge-absent segments were predicted as IDRs, and amongst those that were not, are structurally condensed proteins such as Nup98 and keratins.

Further, to identify that scaffold proteins are enriched in charge-absent regions (Fig 1), we focus on the 51 AA charge-absent segments in proteins without a predicted TM segment. Comparing such proteins with the total number of annotated human proteins in DrLLPS, occurrence with 51 AA charge-absent regions is 26% (scaffold), 8% (regulator), and 4% (client), supporting potential roles in phase separation. Gene Ontology analysis coupled to visualisation with Revigo [51] revealed that proteins containing charge-absent segments are enriched in transcriptional processes and in particular DNA-templated transcription with nuclear localisation (S1 and S2 Figs).

### Charge-absent regions in the human proteome are as hydrophobic as structured proteins

To provide context on the degree of hydrophobicity (Kyte-Doolittle hydropathy, KD, here plotted on a 0 to 1 scale, KD-scaled) of the charge-absent regions, comparison with structured proteins was made. For this purpose, the KD-scaled values were also calculated for each protein in a dataset of 1627 human proteins from PDB structures (non-redundant at 25% sequence identity) [52]. The distribution of KD-scaled values for the charge-absent window distribution is more hydrophobic, on average, than that of the structured protein set (Fig 2). This implies propensity for a condensed phase, perhaps even approaching that of folded proteins, at least in terms of average packing in what is presumably a set of interchanging conformations. These results are biased by selection for the absence of charged amino acids, and it is instructive to also examine the compositions of other amino acids, relative to the structured protein set (Fig 3). Overall, the charge-absent regions are depleted in AAs with aromatic and large hydrophobic sidechains, and are enriched in smaller AAs, in particular Ala, Gly, Pro and Ser, which combine to make up 54% of the AA composition of these 51 residue segments (Fig 3). Calculation of the Shannon entropy for complexity of AA composition [42] revealed that the charge-absent segments are predominantly low complexity regions (LCRs, average entropy 2.9) compared with structured proteins (average 4.0), although the absence of charged AAs will contribute to this result.

**Fig 2.**
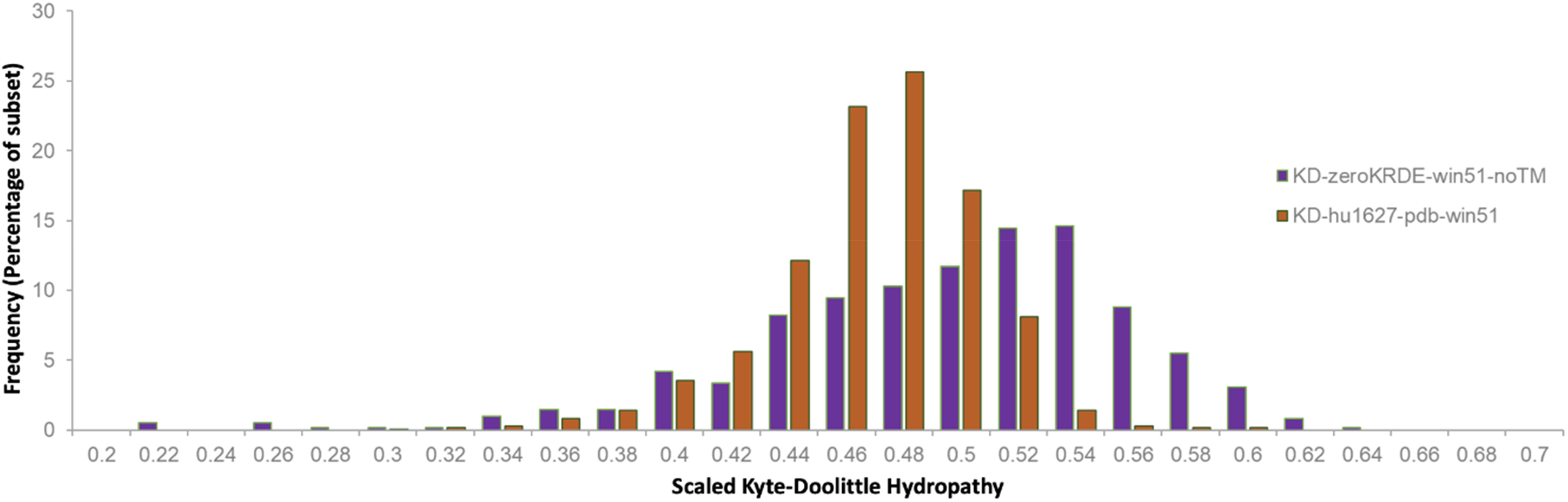
Charge-absent windows have high Kyte-Doolittle hydropathy. Scaled Kyte-Doolittle hydropathy values are shown in histograms for the 623 subset of charge-absent windows of more than 51 AA length (the first occurring such window in a protein of the human proteome), in purple, and for the human 1627 PDB set (orange).

**Fig 3.**
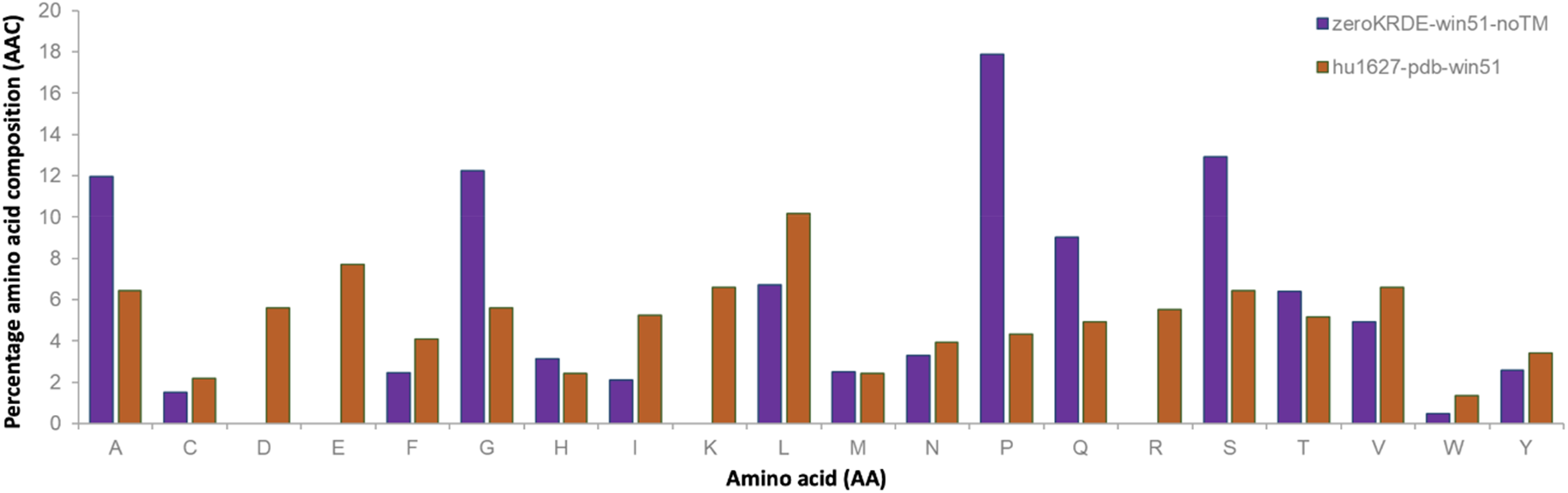
Charge-absent regions are enriched in small amino acids, with a balance of small over large hydrophobics. Percentage AA compositions are compared for the set of charge-absent windows (more than 51 AA, the first such occurrence in each human protein, without predicted TM), in purple, and the 1627 PDB set of structured human proteins (orange).

### Charge-absent regions clustered by amino acid composition

Having established that charge-absent regions in the human proteome are enriched in proteins associated with LLPS and that their hydropathy values are consistent with condensation, we next examined clustering by sequence composition and mapping to specific functions, where these are characterised. The AA composition of each 51-residue segment was calculated using the web tool iFeature [45]. IDRs are enriched in LCRs which in some cases promote LLPS. Different combinations of amino acids were examined, alongside potential links according to function and, more specifically, possible involvement in phase separation. The charge-absent sequences were clustered by amino acid composition (not sequence homology) z-score as described in the methods section. Eleven clusters were identified at the specified cut-off (Fig 4).

**Fig 4.**
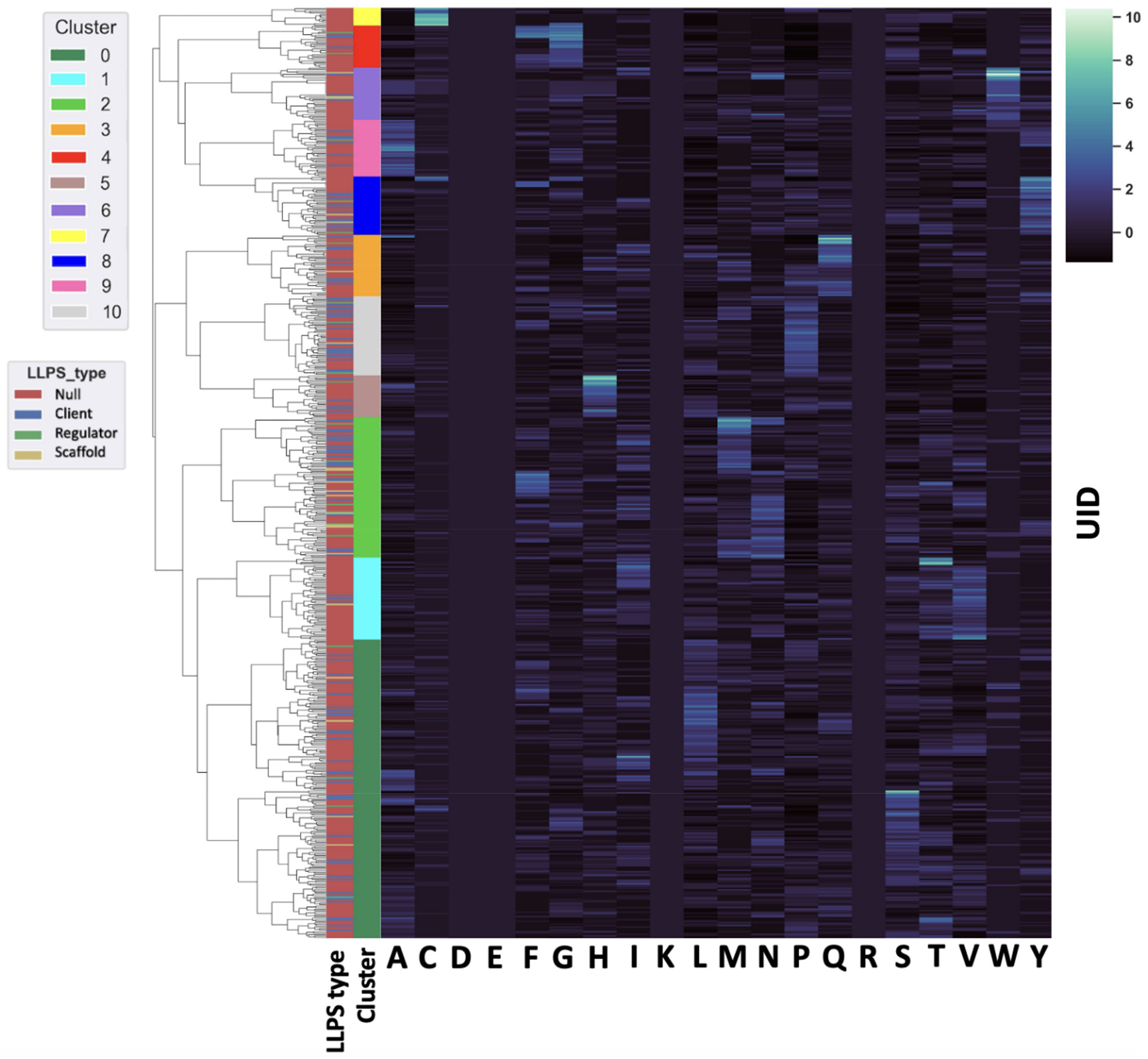
Clustermap of 623 charge-absent 51 AA protein sequences. The sequence segments were normalised by z-score and were clustered by AA composition. Subsets of human proteins are colour coded according to annotation in DrLLPS with Scaffold proteins shown in gold, regulator proteins shown in green and client proteins shown in blue. The null subset covers human proteins without DrLLPS annotation and is coloured in red. The clustermap was also colour-coded by cluster.

Prion-like domains are enriched in uncharged polar amino acids as well as Gly and are often found in RNA binding proteins involved in neurodegenerative diseases caused by protein aggregation such as ALS [61]. Within a list of human proteins with PLDs [47], 68% contained at least one 51 segment absent of charged residues. Proteins with PLDs were found throughout the 11 clusters, apart from Cluster 7. Generally, PLDs have been reported to drive phase separation and, alternatively, to modulate protein phase behaviour [62]. Table 1 gives examples of proteins in each of the clusters, along with amino acid enrichment associated with the cluster, and numbers that have been associated with PLDs. References discussing biological background are given for each example. Some examples are highlighted in the following text, with specific context for charge-absent regions. From combining annotation of a protein as involved in LLPS, with at least one relatively long (> 50 AA) charge-absent region, it does not follow that a charge-absent region necessarily has a central role in phase separation. Even where further characterisation has been reported, this may not be the complete picture. However, there are several examples where charge-absent regions have been directly implicated in LLPS.

**Table 1.** Examples of proteins containing at least one 51 amino acid charge absent region from each cluster. In some of the clusters these proteins have been implicated in phase separation either as scaffolds, clients, or regulators.

| Cluster | AA enrichment | Cluster Size | Proteins with Prion-like domains (PLDs) | Examples |
| --- | --- | --- | --- | --- |
| C0 | S, L | 200 | 37 | YAP1[63], UBQLN2[64, 65], RBM14[66] |
| C1 | T, V | 55 | 7 | BRD3[67], BRD4[68, 69], YY1[70] |
| C2 | M, N | 94 | 41 | TDP-43[12, 36, 38, 71], CREBP[72], p300[72], Nup54, Nup58, Nup62[73], Nup98[74, 75], Nup153 |
| C3 | Q | 41 | 20 | TBP[76, 77], ZNF207[78] |
| C4 | G | 28 | 12 | Loricrin[79] |
| C5 | H | 28 | 3 | Nufip2[80] |
| C6 | W | 35 | 6 | TIA1[81], POLR2A[82], LGALS3[83] |
| C7 | C | 12 | None | None |
| C8 | Y | 39 | 22 | FUS[10, 11, 84, 85], EWSR1[84] |
| C9 | A | 38 | 3 | HOXA13[76] |
| C10 | P | 53 | 10 | WASL[86] |

Bromodomain containing protein 4 (BRD4) is part of cluster 1 and is listed as a scaffold protein in DrLLPS. The N-terminus of BRD4 includes two acetyl-lysine binding bromodomains, separated by an intrinsically disordered region that includes the identified 51 AA segment without charge (232-283) [69]. BRD4 is involved in gene transcription, DNA replication and repair and a short isoform (BRD4S) forms condensates within the cell nuclei [68, 69]. LLPS of BRD4 in the nucleus is mediated by binding of its IDRs and bromodomains to acetylated chromatin and DNA [69], and is diminished upon phosphorylation.

Nucleoporins (Nups) are a family of around 30 intrinsically disordered proteins that make up the nuclear pore complex (NPC). They are in cluster 2 and are known for being rich in Phe and Gly residues, rather than in Met and Asn that is a more common feature of cluster 2 (Table 1). This also indicates that although aromatic AAs are somewhat depleted across charge-absent regions, there are specific families that buck this trend. Nup54, Nup58, Nup62, Nup98 and Nup153 fall in cluster 2. Nups contain multiple Phe/Gly-rich repeats of different kinds throughout their sequence [74] that can promote phase separation through interrepeat interactions [75, 87], and control the transportation of cargo in and out of the nucleus [74]. Since the Phe/Gly-rich regions typically coincide with charge-absent segments, a connection between phase separation and lack of charge can be inferred.

Cluster 2 also includes transcription factors such as TDP-43, CREB-binding protein (CREBBP or CBP), p300, and Forkhead box proteins 1 and 2 (FOXA1, FOXA2). TDP-43 has a methionine-rich charge-absent region (centred on 294-345) that overlaps with the conserved helical region (316-346) known to promote LLPS of TDP-43 [12, 36].

Within the glutamine-rich proteins contained in cluster 3, TATA-box binding protein (TBP) is a transcription factor regulating RNA Pol II activity, with a Gln-rich charge-absent region (including 58 to 109), coincident with an IDR that is reported to drive phase separation. Repeat expansions of the polyQ tail can alter the phase separation capacity of TBP, which in turn can contribute to disease pathogenesis [76, 77]. TBP regulates RNA Pol II transcription of eukaryotes. Moreover, T-cell restriction intracellular antigen 1 (TIA1) is an RNA binding protein residing in cluster 6 that includes overlap of a charge-absent region and a Pro-rich LCD that mediates LLPS, and in which Pro to Leu mutations alter droplet morphology and are disease-associated [81].

It is known that phase separation can also be mediated by combination of charge-absent and other regions, as in the case of FUS, an RNA binding protein in cluster 8. The charge-absent region is located in a PLD rich in Gly, Ser, and Tyr [88, 89]. Tyrosines in the PLD can assist phase separation through cation-π interactions with Arg residues from neighbouring RNA-binding domain of FUS [84]. In addition, Gln residues in FUS also contribute to phase separation, through hydrogen bonding interactions [85]. Furthermore, EWS RNA binding protein 1 (EWSR1) is part of the FUS family of proteins and has similar characteristics to FUS, with a charge-absent region coincident with a PLD. Similar to FUS, EWSR1 phase separates when the PLD interacts with the RBD through cation-π interactions [84]. Collectively, these examples demonstrate that charge-absent regions play a role in phase separation either through homomeric or heteromeric interactions, possibly with charged regions (for example with cation-π interactions).

### Charge modulation of charge-absent regions

It is hypothesised that charge-absent regions (or at least a subset) may be associated with phase separation, by virtue of the absence of desolvation penalty. In this case, the core of some condensates would be devoid of net charge through absence of interacting charges, as opposed for example to those where RNA and RNA-binding proteins combine. They would therefore involve either homomeric or heteromeric interactions, but in either case partner proteins should also contain charge-absent regions. As noted previously, it is also possible that charge-absent regions could supply aromatic groups for partnering with basic residues in cation-π interactions. If charge-absent segments do encourage condensation, then two factors that could alter charge and thus modulate phase separation are phosphorylation and His protonation (at mild acidic pH).

For the 623 human protein subset, the numbers of phosphorylation sites recorded in UniProt that locate to any region within or without a charge-absent 51 AA segment were calculated. Of 2898 total phosphorylation sites from UniProt, only 9 (0.3%) are within 51 AA long charge-absent regions, leaving 99.7% without. For reference, the split of all AAs between within and without charge-absent segments is 4% and 96%, respectively. It is apparent that phosphorylation is depleted in charge-absent regions, despite the propensity in general for phosphorylation within IDRs [90]. Combined Ser and Thr residues are split at 6% within charge-absent regions and 94% without, similar to overall AA numbers, so there is no lack of phosphorylation targets in charge-absent regions. There may however be a lack of defined motifs for protein kinases, since these often involve the charged amino acids that are absent [91]. Whatever the reason, it appears that there exists limited scope for modulation of charge-absent region function by phosphorylation.

Although the normal His sidechain pKa is around 6.3, it is possible that mild acidic pH could lead to protonation (full or partial) and thus exert some influence on charge-absent regions. Percentages of His within and without charge-absent regions (51 AA windows) of the 623 protein subset are 4% and 96%, respectively, matching the overall amino acid split. It is therefore possible that His protonation could be a modulating factor for charge-absent region function, depending on the existence of a mild acidic pH environment. Although the overall His content in charge-absent regions is not enriched, motifs of > 5 consecutive His are more prevalent (11 in 623, versus 70 in the entire human proteome). His tracts with metal ion coordination ability are present in some human transcription factors [92], and separately metal ion-induced condensation has been demonstrated for proteins with engineered hexaHis tags [93]. It is not clear at this stage whether metal ion-dependent association is important for the function of some proteins in the charge-absent region subset, or if it were, how that might couple to the central property (absence of charged residues). One issue is the location of poly-His tracts, for example in BRD4, a hexa-His motif lies at one end of a charge-absent region, bordering a highly charged neighbouring segment. More generally His-rich domains (HRDs) have been associated with targeting of proteins and, as part of LCDs, LLPS activity [94]. An HRD in the YY1 transcription factor, within a region of 33 AAs lacking charged residues, contributes to LLPS and gene expression [70]. The HRD of YY1 is bounded by regions rich in negative charge, raising the possibility of synergistic charge interactions between the neighbouring domains.

## Conclusion

This study focused on protein regions absent of charge, predicted to be largely disordered, and representing a single point in the diagram of fractional positive versus fractional negative charge [19]. Human proteins annotated as scaffold proteins in protein condensates are enriched in charge-absent regions. It is unclear whether charge-absent regions directly drive phase separation, as might be hypothesised from the reduced desolvation energy required, or whether they interact with other regions to promote LLPS. We speculate that charge-absent regions have a propensity to phase separate into relatively compact, but presumably dynamic, structures due to an amino acid composition that in many cases has Kyte-Doolittle hydropathy comparable to that of folded and structured proteins, and without the energetic cost for partially desolvating charged groups. Charge-absent regions are enriched in small hydrophobic AAs and depleted in some hydrophobic AAs with larger sidechains. Amino acid repeats are relatively common in charge-absent regions, compared with charge-containing windows (S3 Fig). We propose that the lack of charge is not solely a result of enrichment of certain amino acids and repeats but is (at least in some cases) key for function. This is supported by the well-characterised example from our charge-absent region dataset, TDP-43. Homomeric interactions of a partly helical segment, that overlaps the charge-absent region, modulate phase separation. Importantly, single site mutations that either alter helical propensity or add charge are associated with disease [36, 38]. It is suggested that charge-absent regions can contribute to the formation of protein condensates by virtue simply of a lack of desolvation penalty for groups bearing net charge. Many differently charged biological systems are known to undergo LLPS, including those with positively and negatively charged domains from within one molecule or between molecules. Condensation with charge-absent regions, where it occurs, would therefore represent just a small subset of phase separating systems, possibly with particular properties in terms of packing (close since there is no charge desolvation) and interchanging conformations (since they are enriched for AAs with small hydrophobic sidechains). Predicting whether a protein will undergo phase separation from sequence is challenging due to the complexity associated with LLPS such as recruitment of other molecules, including ions, proteins, and nucleic acids. While many studies focus on the implications of charge for LLPS, this study reports on regions bearing no charge. The hypothesised role for some charge-absent regions can be investigated by experiments that monitor condensation when charge is added, through mutation, via His protonation and pH-dependence, or with phosphorylation.

## Acknowledgments

The authors would like to acknowledge the University of Manchester and Agency for Science, Technology and Research (A*STAR) for funding as well as the National Supercomputing Centre (NSCC), Singapore for computing facilities.

## Supporting information

**S1 Fig.**
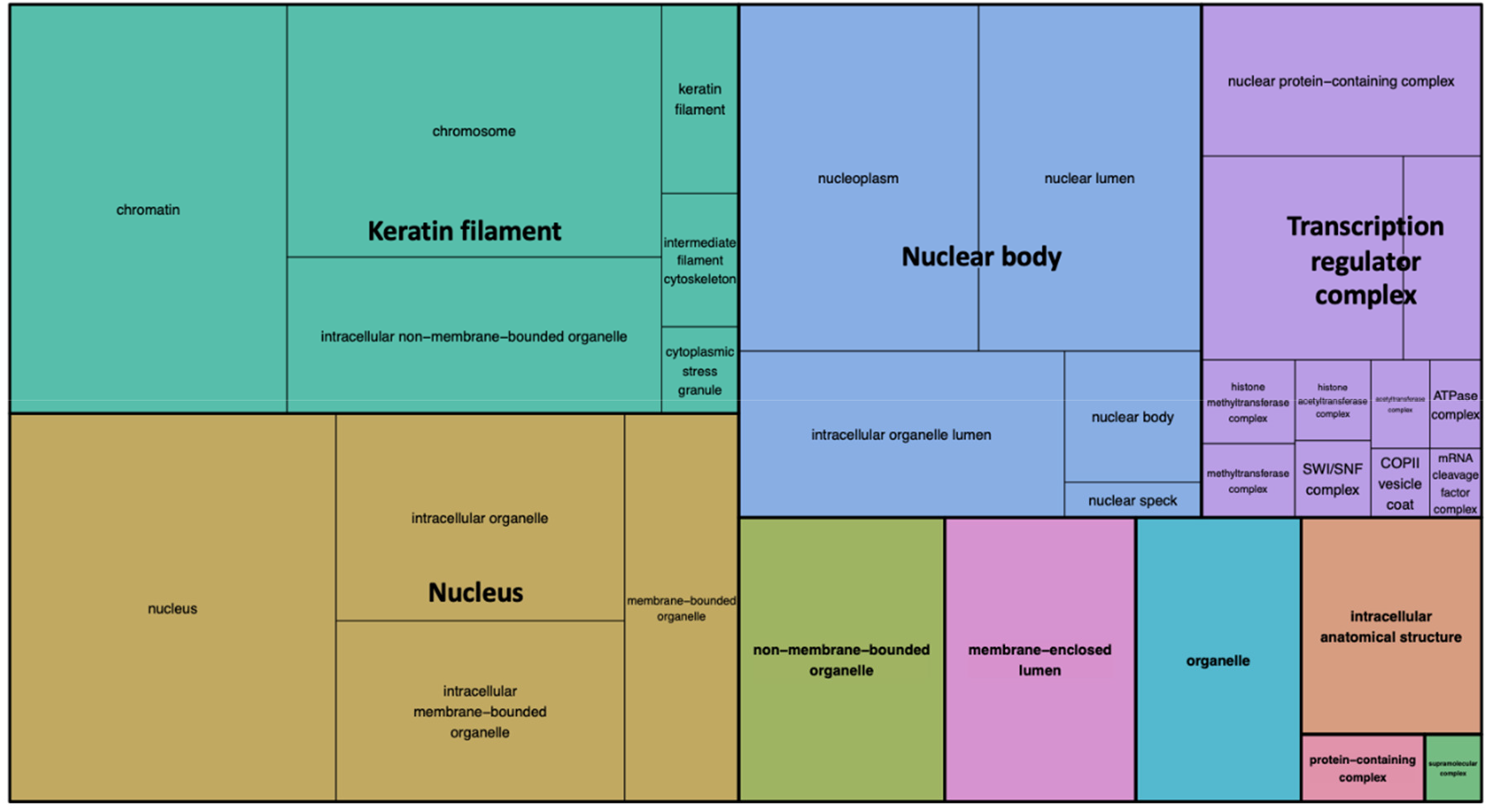
Treemap summarising GO analysis based on cellular component. The analysis was carried out for the subset of proteins containing charge-absent regions.

**S2 Fig.**
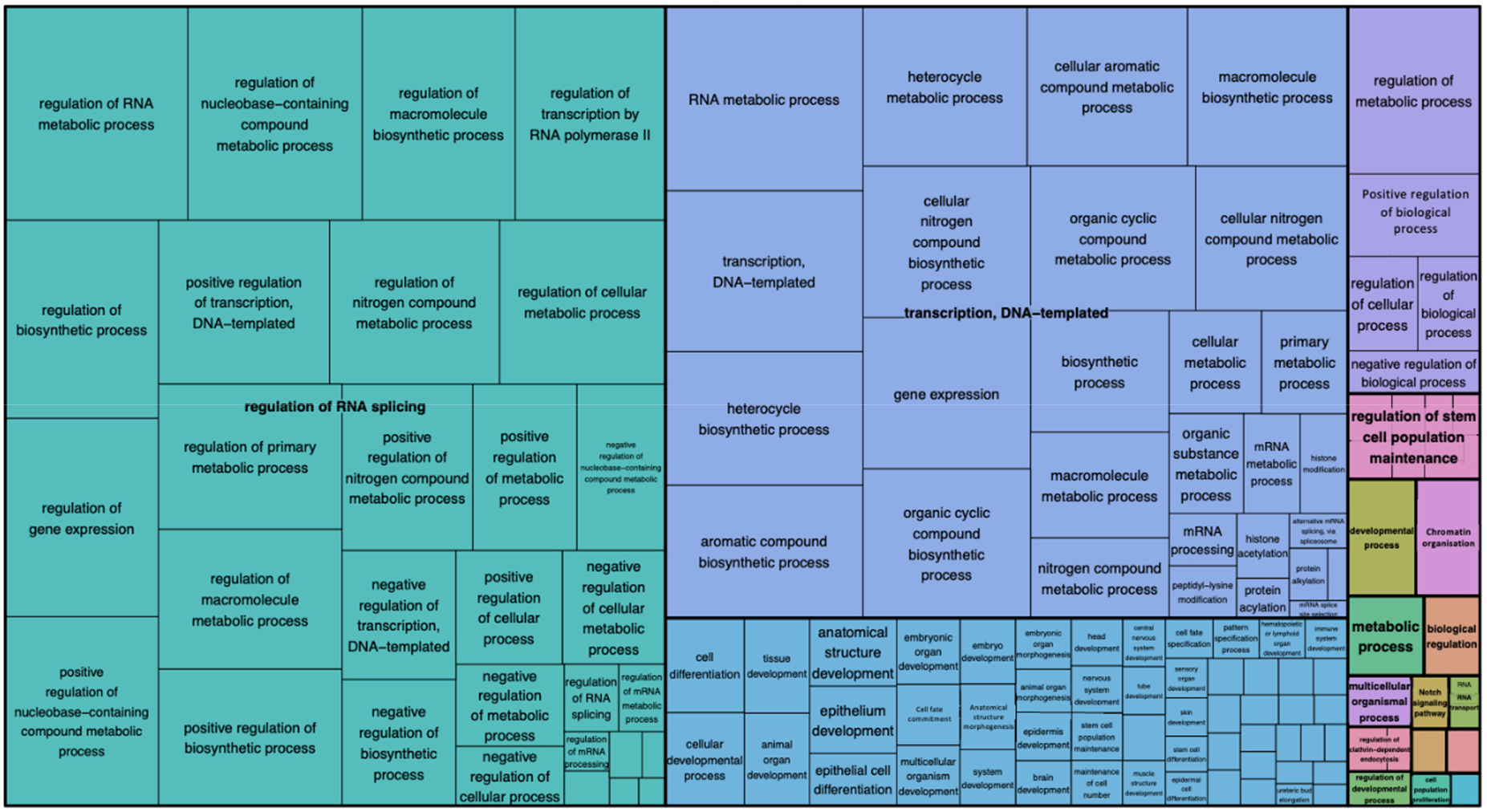
Treemap summarising GO analysis based on biological process. The analysis was carried out for the subset of proteins containing charge-absent regions.

**S3 Fig.**
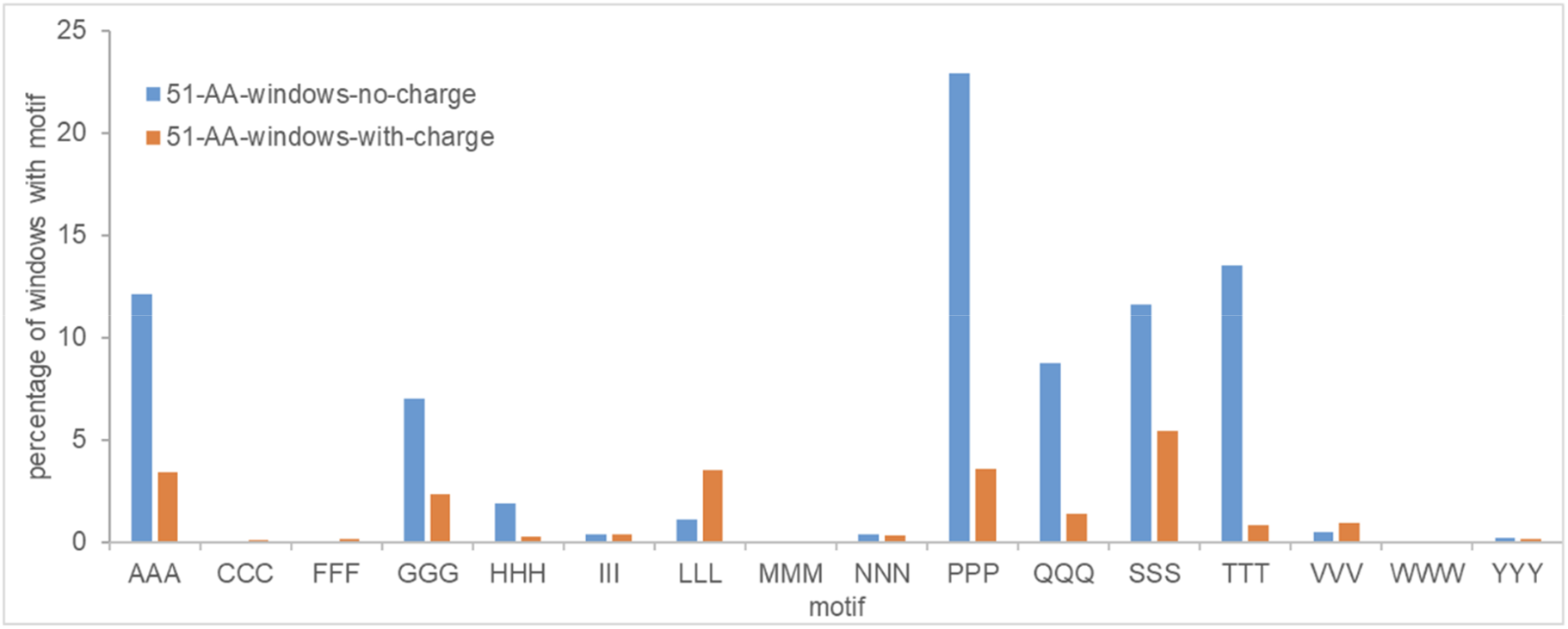
Repeats of certain amino acids are enriched in charge-absent regions. All 51 AA windows in the human proteome (excluding proteins predicted to contain a TM segment) are classified as charge-absent (no-charge, blue) or not (with-charge, orange). The percentage of windows containing a tri-AA motif (excluding Lys, Arg, Asp, Glu) is calculated for the two subsets (no-charge, with-charge).

